# The sensitivity of third party punishment to the framing effect and its brain mechanism

**DOI:** 10.1101/2021.01.11.426181

**Authors:** Jiamiao Yang, Jie Liu, Ruolei Gu, Kexin Deng, Xiaoxuan Huang, Fang Cui

## Abstract

People as third-party observers, without direct self-interest, may punish norm violators to maintain social norms. However, third-party judgment and the follow-up punishment might be susceptible to the way we frame (i.e., verbally describe) a norm violation. We conducted a behavioral and a neuroimaging experiment to investigate the above phenomenon, which we call “third-party framing effect.” In these experiments, participants observed an anonymous player A decided whether to retain her/his economic benefit while exposing player B to a risk of physical pain (described as “harming others” in one condition and “not helping others” in the other condition), then they had a chance to punish player A at their own cost. Participants were more willing to execute third-party punishment under the *harm* frame compared to the *help* frame, manifesting as a framing effect. Self-reported moral outrage toward player A mediated the relationship between empathy toward player B and the framing effect size. Correspondingly, the insula (possibly related to empathy) and cerebellum (possibly related to anger) were activated more strongly under the *harm* frame than the *help* frame. Functional connectivity between these regions showed strongest weight when predicting the framing effect size. These findings shed light on the psychological and neural mechanisms of the third-party framing effect.

**Graphic abstract:** 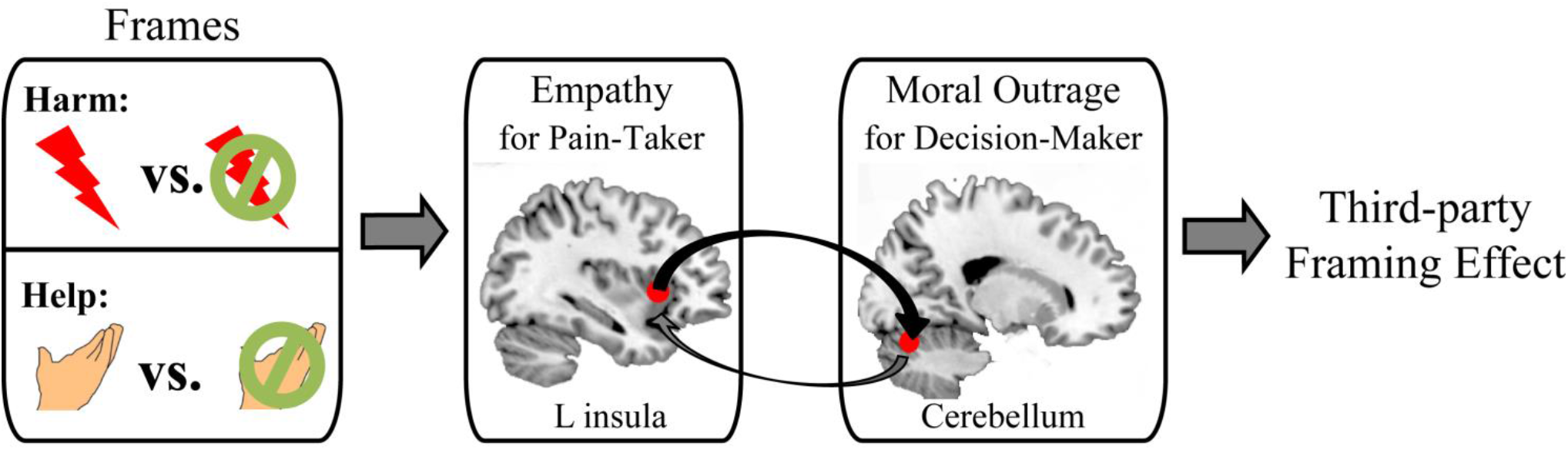

## Introduction

Protests organized by feminists were once often described as “catfights” by American media in the 1960s; as the gender equality movement develops, similar events have been reporting in more favorable words over time (Ashley & Olson, 1998). The above example showcases an attempt to manipulate third-party judgment with framing techniques, which is prevalent in our daily life (e.g., in lawsuits, journalism, and public relations). People make moral judgments on social norm violations even though they are uninvolved, and may then interrupt by punishing norm violators and helping victims (to the extent that they are willing to sacrifice their own self-interest: see Fehr & Fischbacher, 2004; Lotz et al., 2011). However, the judgment about whether something is ethical or morally justifiable from a third-party perspective could be significantly biased by framing techniques (e.g., Entman, 2007). Regarding the value of third-party punishment in maintaining a society (Bendor & Swistak, 2001; Buckholtz & Marois, 2012), investigating its susceptibility to different frames (i.e., verbal statements) and the associated neural underpinnings would be meaningful.

Generally speaking, “framing effect” refers to the phenomenon that our judgment of an object or issue is affected by its description (Tversky & Kahneman, 1981). As pointed out by De Martino et al. (2006), the framing effect could be interpreted as an affect heuristic underwritten by the emotional system. That is to say, positive and negative frames regulate individual preference by evoking emotional responses (Cassotti et al., 2012). It is therefore not surprising that third-party punishment is sensitive to the framing effect, seeing that assessing responsibility and determining an appropriate punishment are both largely based on emotional reactions (Buckholtz et al., 2008). According to the literature, third-party punishment is mainly driven by: (1) empathic feelings to (potential) victims (Dimitroff et al., 2020; Hechler & Kessler, 2018), and (2) moral outrage to norm violators (Nelissen & Zeelenberg, 2009; Salerno & Peter-Hagene, 2013; Treadway et al., 2014). At the brain level, previous studies have shown that empathizing with other’s suffering engages the bilateral insula, anterior cingulate cortex (ACC), and mid-cingulate cortex (MCC) (Fan et al., 2011; Lamm et al., 2011), while moral outrage toward wrongdoing engages the amygdala, hippocampus, cingulate cortex, and basal ganglia (Fumagalli & Priori, 2012). Recently, the cerebellum is also suggested to be involved in the processing of moral-related emotions including anger and disgust (Baumann & Mattingley, 2012; Demirtas-Tatlidede & Schmahmann, 2013; Turner et al., 2007). The aforementioned brain networks have been consistently activated in experimental paradigms of third-party punishment (Krueger & Hoffman, 2016; Stallen et al., 2018).

Accordingly, we investigate the changes of these brain networks in response to the framing effect in two (behavioral and brain-imaging) experiments. Specifically, participants observed an anonymous player made a trade-off between income maximization and helping other persons to avoid a painful shock. Choosing economic benefit was described as either a “harm” to, or “not helping,” other persons in two frame conditions. Then participants could decide whether to punish that player from a third-party perspective. We hypothesized that participants would make more costly punishments in the *harm* frame condition than in the *help frame* condition (see also Liu et al., 2020), regarding that “not harming others” is a stronger moral norm than “helping others” (Crockett et al., 2014). Correspondingly, the brain regions involved in empathic response and/or moral outrage should activate more strongly in the *harm* frame condition. We also examined the relationship between behavioral data and functional connectivity of these regions across frame conditions, so as to improve the knowledge about how third-party punishment is modulated by the framing effect.

## Experiment 1 (behavioral)

### Methods

#### Participants

101 right-handed students were recruited from Shenzhen University to join in Experiment 1. Three of them were excluded due to failures in data recording, leaving 98 participants in the final sample (46 males, age: 21.14 ±1.41 years [mean ± standard deviation]). The study was conducted according to the ethical guidelines and principles of the Declaration of Helsinki and was approved by the Medical Ethical Committee of Shenzhen University Medical School. Informed consent was obtained from all participants prior to the formal experiment.

#### Experimental design and procedures

Our task design combined the “social framing” task with third-party punishment. The task was developed and successfully applied in our recent studies (Gu et al., 2019; Liu et al., 2020; see also Aupperle et al., 2011; Crockett et al., 2014; FeldmanHall et al., 2015). Specifically, one participant plays as the decision-maker (player A) and her/his partner as the pain-taker (player B). In each trial, player A needs to make a decision by choosing between two possible outcomes: loses 5 monetary units (MUs) from her/his own payoff while player B avoids a painful electric shock (i.e., costly helping), or keeps all the payoff while player B receives that shock. Player A determines the occurrence probability (10%, 30%, 50%, 70%, or 90%) of these two outcomes by moving an avatar on a 5-point scale. The two outcomes are described in different ways in two conditions: in the *help frame* condition, the costly helping outcome was described as “help the other person to avoid a shock,” while the other outcome was described as “do not help the other person;” in the *harm frame* condition, the costly helping outcome was described as “give the other person a shock,” while the other outcome was described as “do not give any shock.”

In this study, all the participants acted as a third-party observer (player C) and received 5 MUs in each trial of the task. After they observed each of player A’s choice, participants were given a chance to punish player A by spending their own 5 MUs, such that each MU spent would reduce 2 MUs from player A’s final payoff (Fig. 1A). Before the formal task, participants were told that the interactions between players A and B were recorded in a previous study, but player A had not yet received her/his payoff. They were also told that the MUs during the task would be converted to real monetary bonus. Thus, their follow-up decisions would determine not only their own payoff (fixed baseline payment + bonus) but also that of player A’s. In reality, both players A and B were putative players (see also Wang et al., 2017). Player A’s choices were pseudo-randomly set up such that the number of trials in each condition was the same.

**Figure 1.**
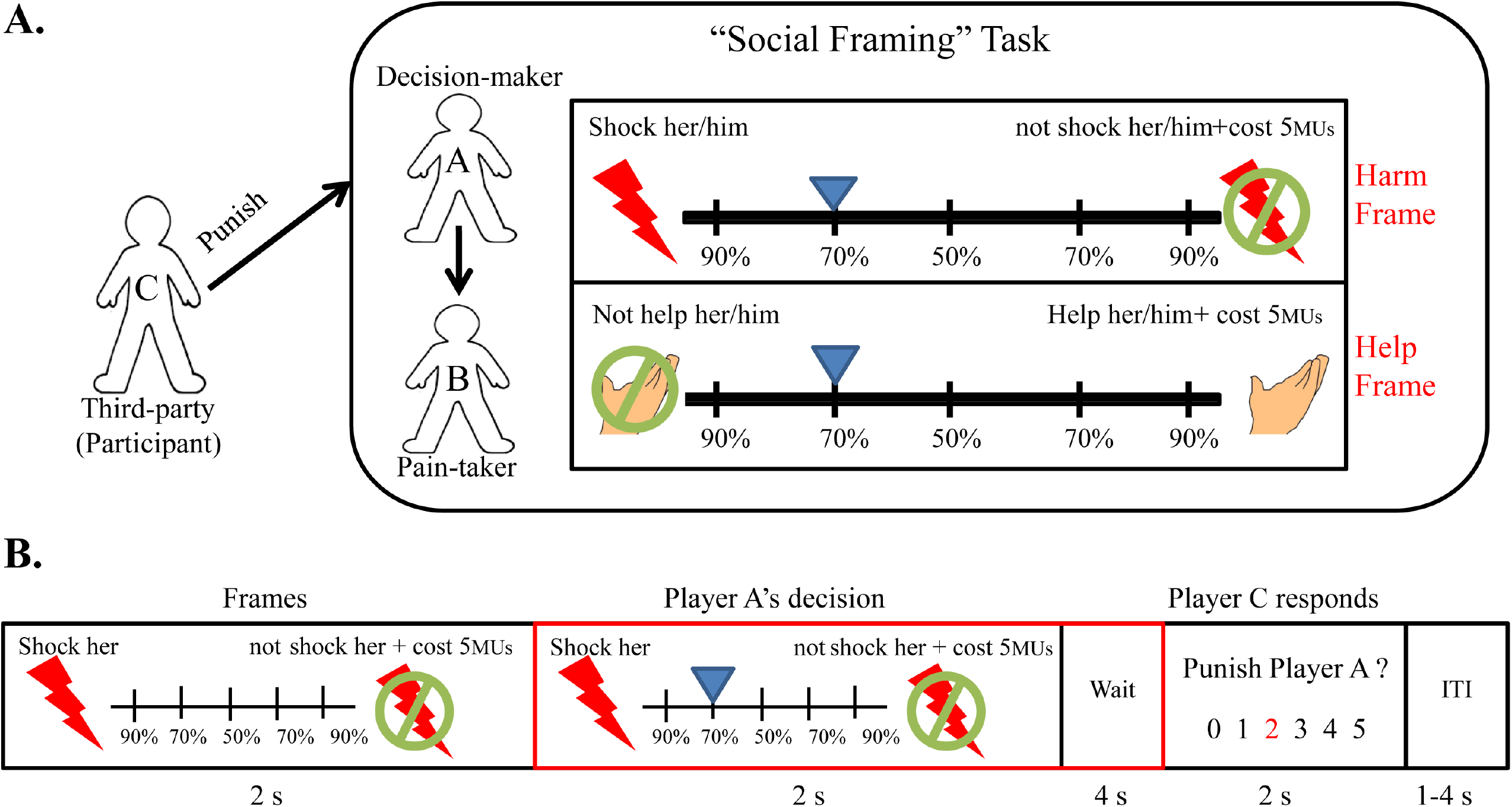
Task design and experimental procedure. (A) The real participant (player C) observed a “social framing” task in which player A decided the probability for player B to receive a painful electrical shock. Then the participant could decide whether and how to punish player A with her/his own income. (B) Schematic of an example trial in Experiment 2. The event for brain-imaging analysis is marked with a red rectangle.

The present experiment employed a 2 (*frame*: harm versus help) × 5 (*moral level of player A’s choice* [“*moral level*” for short]) within-subject design. Here, the five levels of the *moral level* factor were: highly pro-helping (90% probability of costing own money to save player B from a shock), medium pro-helping (70% probability of costing own money), neutral (50%), medium non-helping (70% probability of keeping own money), and highly non-helping (90% probability of keeping own money), respectively. Each condition (2 × 5) repeated twice, resulting in 20 trials through the task.

In each trial, the real participant (player C) first observed two possible outcomes (which were presented in different ways in two frame conditions) for 2 s. The position of each outcome (left/right) was counterbalanced across trials. After that, a blue avatar moved on a 5-point scale to show player A’s decision process for 2s. Each participant then input numbers with numerical keys on a keyboard to answer three questions: (1) *altruistic punishment*: “how many MUs you would like to pay for punishing player A?” (0~5 MUs); (2) *empathic feeling for player B*: “how unpleasant player B would feel?” (1: not unpleasant at all ~ 5: extremely unpleasant); (3) *moral outrage against player A*: “how angry do you feel about player A? (1: not angry at all ~ 5: extremely angry). The order of these three questions was counterbalanced across participants. The whole task lasted for approximately 5 mins.

### Results

As mentioned in the Introduction, we focused on whether different frames would bias participants’ judgment about player A’s choice and then modulate their altruistic punishment (i.e., third-party framing effect).

First, to examine whether the strength of altruistic punishment was sensitive to the two within-subject factors (i.e., *frame* and *moral level of player A’s choice*), we performed a 2 × 5 repeated-measures analysis of variance (ANOVA) on punishment data (in MUs). The results showed significant main effects of both factors (*frame*: *F*_(1, 97)_ = 35.64, *p* < 0.001, *η_p_^2^* = 0.27; *moral level: F*_(4, 388)_ = 164.38, *p* < 0.001, *η_p_^2^* = 0.63) (*ps* < 0.001) as well as an interaction (*F*_(4, 388)_ = 10.15, *p* < 0.001, *ηp^2^* = 0.10). Pairwise comparison showed that when player A was highly pro-helping, participants’ punishment was not significantly different between two frames (*harm frame*: 0.09 ± 0.03; *help frame*: 0.08 ± 0.03, *p* = 0.53); in other four conditions (medium pro-helping, neutral, medium non-helping, and highly non-helping), participants punished more under the *harm* frame than the *help* frame (*ps* < 0.015) (Fig. 2A).

**Figure 2.**
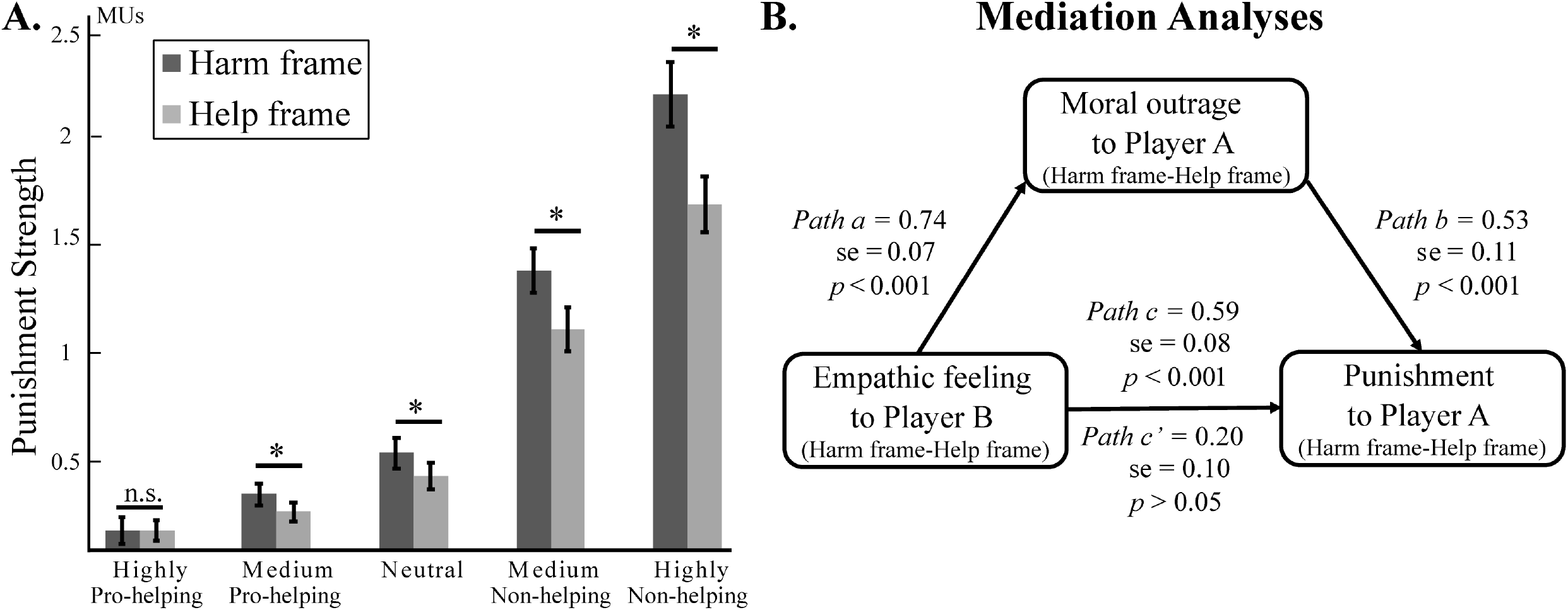
Behavioral results of Experiment 1. (A) The strength of third-party punishment for each moral level in the *harm* frame and *help* frame conditions. (B) Results of the mediation analyses.

We then calculated the difference of participants’ punishment between the *harm* frame and the *help* frame as an index of third-party framing effect size. Pearson’s correlation analysis revealed that the framing effect sizes in different conditions were significantly correlated with each other (*ps* < 0.002) except when player A was highly pro-helping (*ps* > 0.126).

Overall, these results revealed a significant third-party framing effect (i.e., altruistic punishment was increased under the *harm* frame than the *help* frame) except when player A was highly pro-helping. In the other four conditions (i.e., medium pro-helping, neutral, medium non-helping, and highly non-helping), the third-party framing effect was homogenous. We, therefore, averaged each participant’s framing effect size in these four conditions as a general behavioral index. Also, we calculated the difference in “empathic feeling for player B” as well as “moral outrage against player A” between the two frames, in the same way as we calculated the third-party framing effect (i.e., *harm* frame minus *help* frame). One sample *t*-tests revealed that these three indexes were all significantly larger than zero (framing effect size: 0.23 ± 0.04, *t*_(97)_ = 6.13, *p* < 0.001; empathic feeling: 0.28 ± 0.04, *t*_(97)_ = 7.07, *p* < 0.001; moral outrage: 0.32 ± 0.04, *t*_(97)_ = 8.43, *p* < 0.001). Pearson’s correlation analysis revealed that these three indexes were all significantly correlated with one another (*rs* > 0.63, *ps* < 0.001).

We then run a mediation analysis with the difference in empathic feeling as X, the difference in moral outrage as M, and the third-party framing effect as Y. This analysis revealed a significant full mediation effect of M between X and Y. Normal theory tests showed a significant a path (*t*_(97)_ = 11.30, *p* < 0.001) and a significant b path (*t*_(97)_ = 5.06, *p* < 0.001). A significant c path (*t*_(97)_ = 7.86, *p* < 0.001) became marginally significant (*t*_(97)_ = 1.94, *p* = 0.054) when the mediator was adjusted. The bias-corrected confidence interval was between 0.21 and 0.61. The mediation effect accounted for about 66.30% of the total effect (Fig. 2B).

## Experiment 2 (fMRI)

### Methods

#### Participants

To determine an appropriate sample size, a priori power analysis was conducted using the G*Power 3.1 (Faul, Erdfelder, Lang, & Buchner, 2007). This analysis revealed that 29 participants were required to reach a good statistical power of 0.85 to detect median sized (f = 0.25) effects with an alpha value of 0.05 for a 2 × 5 within-subjects analysis of variance (ANOVA). To account for possible dropouts or errors during the experiment thirty-five right-handed participants who did not participate in Experiment 1 were recruited from Shenzhen University to join in the fMRI experiment. Four of them who had excessive head movements > 2° in rotation or > 2 mm in translation during the scanning were excluded, leaving 31 participants in the final sample (14 females, age: 20.62 ± 1.96 years [mean ± standard deviation]). The study was conducted according to the ethical guidelines and principles of the Declaration of Helsinki and was approved by the Medical Ethical Committee of Shenzhen University Medical School. Informed consent was obtained from all participants before the formal experiment.

#### Experimental design and procedures

The task design was the same as in Experiment 1, but the experimental setting was adjusted for fMRI scanning. Before the scanning, participants were familiarized with the task with a practice block consisting of 8 trials. In each trial, the real participant (player C) first observed two possible outcomes in different frames for 2 s. After that, a blue avatar moved on a 5-point scale to show player A’s decision process for 2 s. Each participant then waited for another 4 s before they could press one of two pre-assigned buttons on an MRI-compatible button-box to indicate how many MUs (0~5) s/he would like to pay for punishing player A. The participant had 2 s to choose the preferred number (turned from black to red), the starting point of which was randomized across trials. Finally, the inter-stimulus interval was set as 1~4 s (Fig 1B). Each condition (2 × 5) contained 24 trials and there were 240 trials in total. The experiment, which consisted of four runs of 60 trials, lasted for approximately 1 h.

#### Neuroimaging data acquisition and preprocessing

We used a Siemens TrioTim 3.0T MRI machine for data acquisition. Functional volumes were acquired using a multiple slice T2-weighted echo planar imaging (EPI) sequences with the following parameters: repetition time = 2000 ms, echo time = 30 ms, flip angle = 90°, field of view = 224 × 224 mm^2^, 33 slices covering the entire brain, slice thickness = 3.5 mm, voxel size = 3.5 × 3.5 × 3.5 mm^3^.

fMRI data were preprocessed in SPM12 (Wellcome Department of Imaging Neurosciences, University College London, U.K., http://www.fil.ion.ucl.ac.uk/spm). Images were slice-time corrected, motion-corrected, and normalized to Montreal Neurological Institute (MNI) space for each individual with a spatial resolution of 3 × 3 × 3 mm^3^. Images were then smoothed using an isotropic 6-mm Gaussian kernel and high-pass filtered at a cutoff of 128 s.

### Neuroimaging data analysis

#### Whole-brain activation analysis

Statistical parametric maps were generated on a voxel-by-voxel basis with a hemodynamic model to estimate brain response. We mainly focused on analyzing participants’ brain signals when they watched the presentation of player A’s choice, which is highlighted with a red rectangle in Fig. 1B. In our general linear model (GLM), the presentation of possible outcomes, player A’s choice, participant(as player C)’s response, as well as interstimulus intervals were included in the GLM at the single-participant level. The six rigid-body parameters were also included in the GLM to exclude head-motion nuisance. The independent regressors which we analyzed in the GLM were different frames (*harm* versus *help*) under different moral levels of player A’s choice (highly pro-helping, medium pro-helping, neutral, medium non-helping, and highly non-helping), namely 10 independent regressors.

Since we are most interested in the third-party framing effect, we combined four moral levels (medium pro-helping, neutral, medium non-helping, and highly non-helping) to focus on the main effect of frames; as mentioned above, the highly pro-helping condition was excluded because no framing effect was found in this condition. For the group-level analysis, we conducted a one-sample *t*-test using the whole brain as the volume of interest to localize the differences in brain activity between the *harm* frame and the *help* frame. Besides, a regression analysis was also conducted to explore which brain areas were activated stronger in the contrast of *harm* minus *help* frame, as a function of the third-party framing effect.

#### Task-dependent FC for predicting the third-party framing effect

Regarding the brain network that responded to the third-party framing effect, we tested whether there was sufficient information in the connectivity patterns within this network to predict individual difference in behavioral decisions. To do this, we first defined the regions of interest (ROIs) as brain areas of which the activation level: (1) was significantly higher in the contrast of *harm* minus *help* frame, and (2) covaried with the behavioral framing effect size. Five brain regions were found to meet the above two criteria (see the Results section). We then estimated the FC between each pair of the ROIs using psychophysiological interactions (PPI). We then drew spheres (radius = 6 mm) at the coordinates of all the five significant peaks of activity (see Table 3) localized in the overlap between the T-contrast map of *harm* > *help* and the regression map that covaried with the framing effect size.

#### Calculation of FC matrix

Task-dependent connectivity was estimated using a linear model equivalent to the general linear model (GLM) typically used in fMRI analysis with several additional regressors. Specifically, there were three kinds of regressors per linear model: one task regressor convolved with a subject-specific hemodynamic response function (equivalent to the regressors in a typical GLM), a contrast of the task-dependent activity, and a task-dependent connectivity regressor for the task condition consisting of each seed region’s time series during which each participant was performing the task.

We calculated whole-brain task-dependent FC maps with all the five ROIs as seed regions and then extracted parameter estimates of each FC map within each ROI, so as to obtain a matrix that represented the FC strength between each pair of ROIs for each participant.

#### Linear relevance vector regression (RVR)

We selected the RVR due to its high prediction performance in brain-behavior/ cognition mapping (Cui & Gong, 2018). RVR is a Bayesian framework for learning sparse regression models, which does not include an algorithm-specific parameter and therefore does not require extra computational resources for parameter estimation (Tipping, 2000). In RVR, only some samples (fewer than the training sample size), called “relevance vectors,” are used to fit the model: y(x) = ∑w_i_ τ_i_ + ∈. Leave-one-out-cross-validation (LOOCV) was used to calculate the prediction accuracy (i.e., Pearson correlation coefficient between the predicted and actual labels). For each round of LOOCV, one participant was designated as the test sample and the remaining participants were used to train the model. The predicted score was then obtained from the feature matrix of the tested sample.

The significance level was computed based on 1000 permutation tests. For each permutation test, the prediction labels (i.e., participant’s third-party framing effect) were randomized, and the same RVR prediction process used for the actual data was carried out. After 1000 permutations, a random distribution of accuracies was obtained and the *p*-value was correspondingly calculated as *p* = (number of permutation tests < actual accuracy +1)/(number of permutation test + 1).

#### Weight-based brain network

The feature weight indicates the importance of each feature in the regression model. For the significant prediction models (permutation *p* < 0.05), the most predictive connections with the highest absolute weights were extracted. In the linear RVR model, connections with positive weights indicate that increased FC predicts a stronger third-party framing effect, while connections with negative weights indicate that increased FC predicts a weaker third-party framing effect.

#### Validation analysis

To determine whether our major findings were sensitive to our choices of correlation thresholds for connectivity, we recomputed the FC with different thresholds (0.1) and then re-performed the machine learning analysis.

#### ROI analysis

As we found in the machine learning-based FC analysis, the FC between the left insula and cerebellum was the strongest predictor of the third-party framing effect. We assumed that the insula represents the empathizing process, while the cerebellum represents moral emotions such as anger and disgust. To examine the above hypotheses, we ran an additional ROI analysis. Specifically, we selected ROIs in the left insula (MNI: [−36 16 2]) from a meta-analysis focusing on empathy (Fan et al., 2011), and selected ROIs in the cerebellum which are associated with the processing of anger (MNI: [18 −72 −24]) and disgust (MNI: [−16 −70 −21]), respectively, according to another study (Baumann & Mattingley, 2012). Besides, to test whether other functions of the cerebellum were also involved in our task, ROIs in the cerebellum representing mentalizing (MNI: [24 −82 −28] and [16 −84 −26]) from a meta-analysis (Van Overwalle et al., 2014) were also selected. Mean parameter-estimates were calculated around the peaks of these ROIs (for each participant and each frame condition). We then performed *t*-tests between the *harm* frame and the *help* frame condition on the extracted values to examine their difference.

#### Dynamic causal modeling (DCM)

In the above FC analysis, the brain connection between the left insula and cerebellum showed the strongest predictive weight in relation to the third-party framing effect (i.e., stronger than all the other connections in the 5 node-network). The direction of this connection was further validated through DCM. We used DCM12 to examine the effective connectivity between the above two brain regions during the task (Friston et al., 2003). The first eigenvariate for the single-participant time courses was extracted from volumes located in the left insula and cerebellum, using a sphere with a 6-mm radius that was centered at individual maxima and adjusted for the effects of interest. The ROI time series were extracted from within the whole-brain activation for the *harm* frame and *help* frame conditions.

To determine the driving input (matrix C) and modulation effect (matrix B), we fixed the intrinsic connection between the two regions as bilateral connections. Our model space consisted of 9 models, which were differentiated by where the modulation effect (matrix B) and the driving input (matrix C) took place. Thus we included all combinations of possible modulations between the left insula and the cerebellum (Figure 5A, unidirectional modulation effect: left insula → cerebellum, cerebellum → left insula; bidirectional modulation effect: left insula ↔ cerebellum).

We then permutated driving input into each node, respectively, or both of the two nodes. We finally specified all 9 models separately for each run and each participant. We then estimated all models and subjected them to the random-effect Bayesian Model Selection to select the best-fitted model from our model space based on the model evidence (Stephan et al., 2010).

## Results

### Behavioral results

We performed a 2 × 5 repeated measures ANOVA on altruistic punishment and found that the main effects of both within-subject factors (*frame*: *F*_(1,30)_ = 28.43, *p* < 0.001, *η_p_^2^* = 0.49; *moral level*: *F*_(4, 120)_ = 187.70, *p* < 0.001, *η_p_^2^* = 0.86) as well their interaction (*F*_(4, 120)_ = 16.43, *p* < 0.001, *ηp^2^* = 0.35) were significant. Pairwise comparison showed that when player A was highly pro-helping, participants’ altruistic punishment was not significantly different between two frames (*harm* frame: 0.18 ± 0.04; *help* frame: 0.19 ± 0.05, *p* = 0.88); meanwhile, in other four conditions (medium pro-helping, neutral, medium non-helping, and highly non-helping), participants punished more under the *harm* frame than the *help* frame (*ps* < 0.008) (Fig. 3A). As in Experiment 1, these results confirmed the third-party framing effect in four conditions. Accordingly, we then used the averaged of these conditions to index each participant’s third-party framing effect size, as we did in Experiment 1(Fig. 3B).

**Figure 3.**
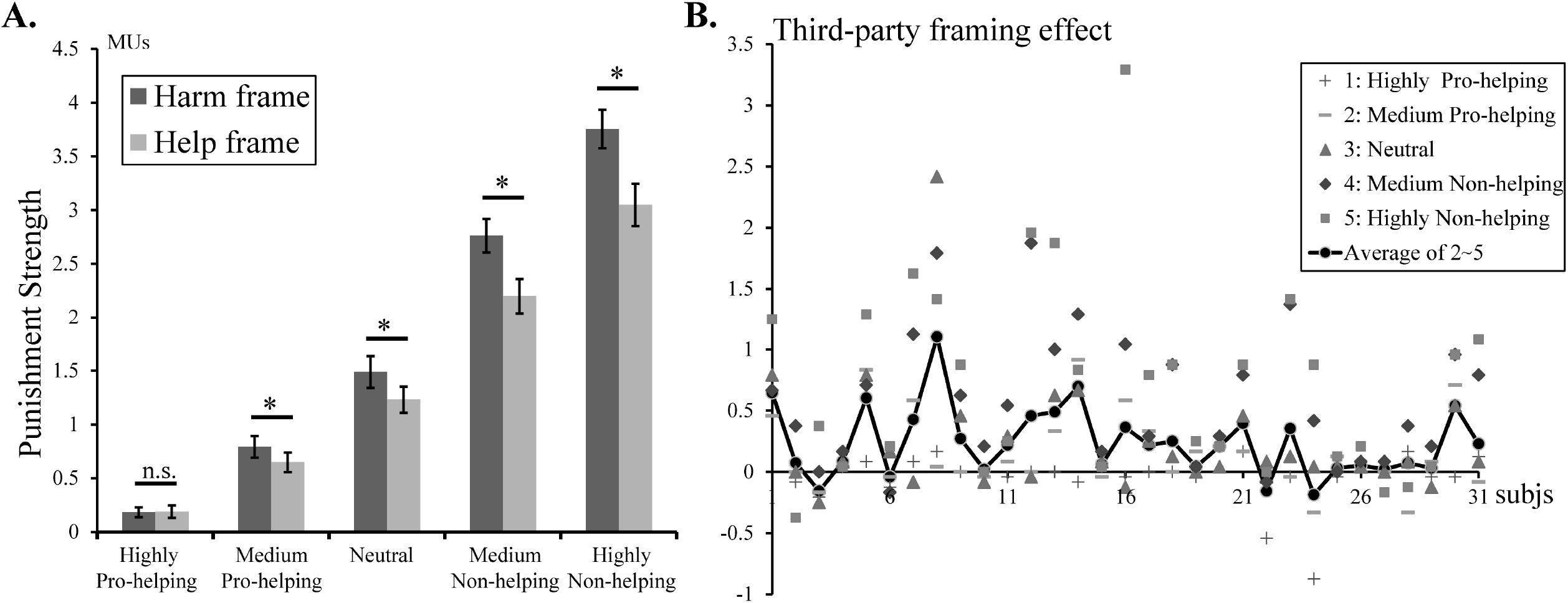
Behavioral results of Experiment 2. (A) The strength of third-party punishment for each moral level in the *harm* frame and *help* frame conditions. (B) The third-party framing effect size for each participant, calculated as the average across four moral levels except “highly pro-helping,” is represented as the solid black line in panel B (*: *p* < 0.01; n.s.: non-significant).

### fMRI results

#### Whole-brain activation analysis

Contrasting the brain activation level under the *harm* frame and the *help* frame, we found significantly stronger activation in the bilateral insula (left peak MNI [−36, 6, 0], right peak MNI [42, 6, −3]), left supplementary motor area (peak MNI [−9, −3, 69]), and left middle frontal gyrus (peak MNI [−42, −21, 45]) (Fig. 4A and Table 1). The regression analysis showed that the activation of the bilateral insula (left peak MNI [−33, 6, 0], right peak MNI [33, 6, 9]) significantly covaried with the third-party framing effect size (Fig. 4B and Table 2). The results of the above two analyses overlapped in five significant clusters, namely the left insula (peak MNI [−36, 6, 0], [−45, 6, 9]), right insula (peak MNI [39, 3, 6]), median cingulate and paracingulate gyri (peak MNI [3, −9, 42]), superior part of the cerebellum (peak MNI [9, −63, −21]) (Fig. 4C and Table 3).

**Figure 4.**
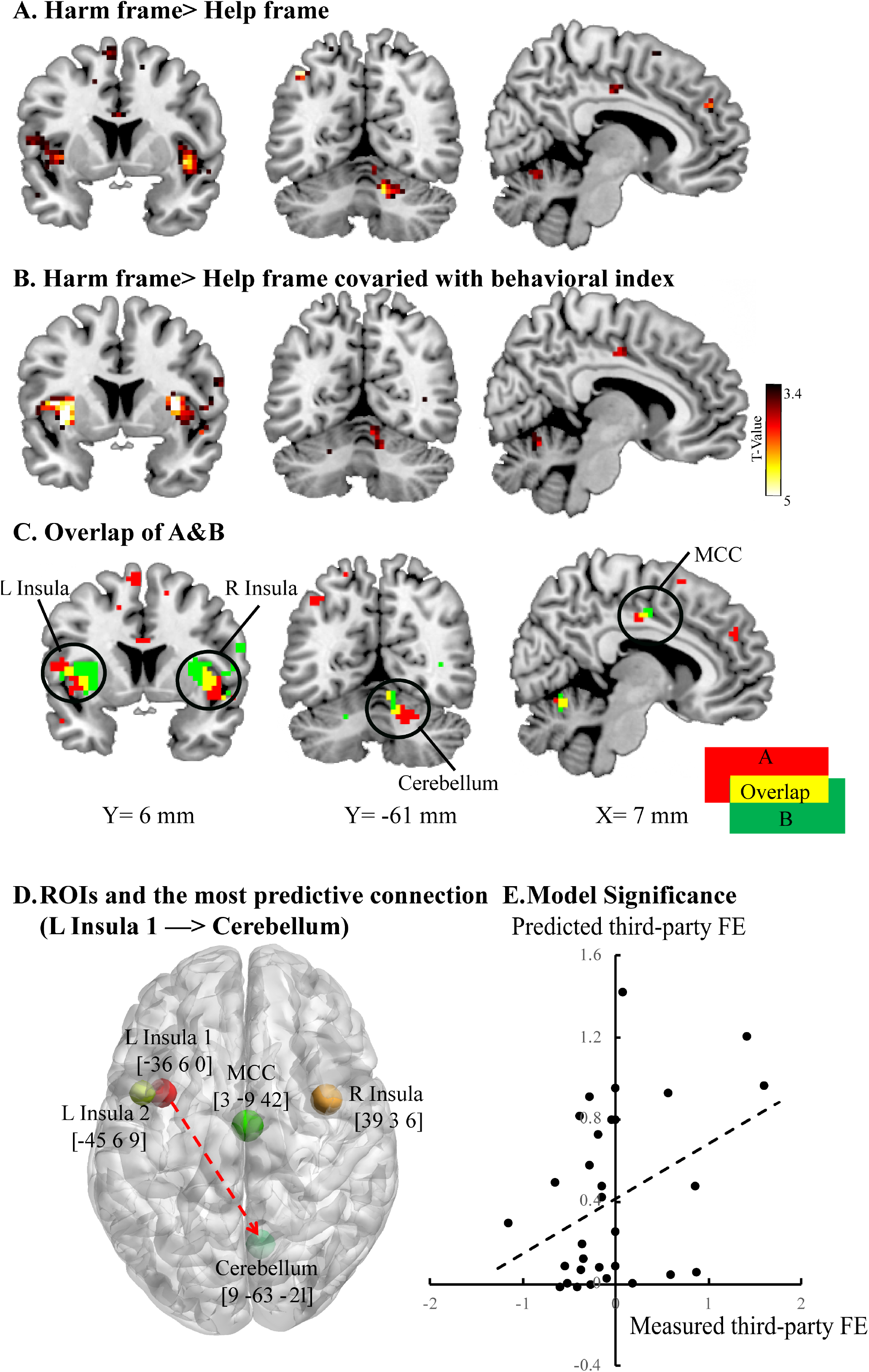
Results of brain activation in Experiment 2. (A) Brain activation of the contrast under the *harm* frame > *help* frame. (B) Contrast of the *harm* frame > *help* frame covaried with individual third-party framing effect at the group level. (C) The overlap between A and B. (D) The 5-node brain network related to the third-party framing effect. (E) Correlation between actual and predicted framing effect (FE) scores within the 5-node brain network.

**Table 1.** Brain activations of the contrast under the *harm* frame > *help* frame.

| Hem. | Brain Region | BA | Coordinates |  |  | Vol. | T |
| --- | --- | --- | --- | --- | --- | --- | --- |
|  |  |  | (X, Y, Z) |  |  |  |  |
| R | Insula* | 48 | 42 | 6 | -3 | 86 | 5.56 |
| L | Supplementary motor area* | 6 | -9 | -3 | 69 | 101 | 5.49 |
|  | Supplementary motor area | 6 | -6 | -12 | 54 |  | 3.95 |
| L | Precentral gyrus | 6 | -27 | -12 | 63 | 41 | 5.4 |
| L | Angular gyrus | 39 | -39 | -63 | 48 | 26 | 4.92 |
| L | Anterior cingulate and paracingulate gyri | 32 | -12 | 45 | 12 | 50 | 4.77 |
|  | Superior frontal gyrus, medial orbital | 10 | -12 | 42 | -6 |  | 3.91 |
| L | Middle frontal gyrus* | 4 | -42 | -21 | 45 | 147 | 4.71 |
|  | Postcentral gyrus | 2 | -51 | -30 | 51 |  | 4.25 |
|  | Inferior parietal | 2 | -42 | -30 | 39 |  | 4.24 |
| L | Insula | 48 | -36 | 6 | 0 | 72 | 4.68 |
|  | Inferior frontal gyrus, opercular part | 46 | -57 | 9 | 12 |  | 4.13 |
|  | Superior temporal gyrus | 21 | -51 | 0 | -9 |  | 4.01 |
| R | Superior frontal gyrus, medial | 10 | 15 | 54 | 9 | 14 | 4.57 |
|  |  |  | -21 | 33 | 30 | 16 | 4.39 |
|  |  |  | 12 | -60 | -27 | 46 | 4.29 |
|  | Cerebellum | 19 | 21 | -63 | -27 |  | 4 |
| L | Middle frontal gyrus | 46 | -27 | 45 | 30 | 36 | 4.28 |
|  | Superior frontal gyrus, dorsolateral | 46 | -24 | 45 | 21 |  | 3.94 |
| L | Median cingulate and paracingulate gyri | 23 | -3 | -24 | 45 | 38 | 4.01 |
|  | Median cingulate and paracingulate gyri | 23 | -3 | -12 | 42 |  | 3.9 |
|  | Median cingulate and paracingulate gyri | 23 | 6 | -15 | 39 |  | 3.69 |
| R | Middle frontal gyrus | 46 | 30 | 45 | 18 | 14 | 3.99 |
| R | Superior frontal gyrus, medial | 32 | 9 | 48 | 30 | 12 | 3.99 |
| L | Middle temporal gyrus | 21 | -60 | -21 | -3 | 37 | 3.95 |
|  | Middle temporal gyrus | 21 | -54 | -18 | -12 |  | 3.84 |

Note: all the results reported above were significant at $p < 0.001$ , $k > 10$ , uncorrected at the voxel level; \* indicates the cluster-level *FWE* correction at $p < 0.05$ .

**Table 2.** Whole-brain activations based on group-level regression analysis predicting the behavioral third-party framing effect.

| Hem. | Brain Region | BA | Coordinates |  |  | Vol. | T |
| --- | --- | --- | --- | --- | --- | --- | --- |
|  |  |  | (X, Y, Z) |  |  |  |  |
| L | Insula* | 48 | -33 | 6 | 0 | 289 | 6.58 |
|  |  | 34 | -30 | 0 | -9 |  | 5.53 |
|  |  | 48 | -36 | -3 | 6 |  | 5.29 |
| R | Insula | 48 | 33 | 6 | 9 | 165 | 6.39 |
|  |  | 48 | 36 | -3 | -6 |  | 4.77 |
|  | Temporal pole: superior temporal gyrus | 38 | 54 | 9 | -9 |  | 4.48 |
|  |  | 21 | -42 | -45 | 9 | 29 | 5.26 |
|  |  | 41 | -36 | -42 | 15 |  | 4.32 |
| L | Cerebellum_Superior | 30 | -21 | -30 | -24 | 19 | 5.2 |
| L | Middle temporal gyrus | 21 | -51 | -6 | -24 | 26 | 5.01 |
| L | Median cingulate and paracingulate gyri |  | -6 | -21 | 45 | 51 | 4.74 |
|  | Median cingulate and paracingulate gyri | 23 | 3 | -9 | 45 |  | 4.21 |
| L | Rolandic operculum | 48 | -42 | -21 | 15 | 46 | 4.72 |
|  | Superior temporal gyrus | 48 | -51 | -27 | 18 |  | 3.69 |
| R | Postcentral gyrus | 22 | 63 | -12 | 15 | 26 | 4.45 |
|  | Rolandic operculum | 48 | 45 | -15 | 15 |  | 3.94 |
|  |  |  | -9 | -27 | 0 | 19 | 4.32 |
|  |  |  | -15 | -39 | 39 | 14 | 4.21 |
| R | Cerebellum_Superior | 18 | 9 | -63 | -21 | 31 | 4.07 |
|  | Cerebellum_Superior | 18 | 9 | -63 | -12 |  | 3.78 |
|  | Cerebellum_Superior | 37 | 18 | -54 | -24 |  | 3.78 |
|  | Paracentral lobule |  | 0 | -24 | 72 | 14 | 4.04 |
| R | Supramarginal gyrus | 48 | 45 | -33 | 27 | 12 | 3.74 |

Note: all the results reported above were significant at $p < 0.001$ , $k > 10$ , uncorrected at the voxel level; \* indicates the cluster-level *FWE* correction at $p < 0.05$ .

**Table 3.** Overlap between the contrast map and the regression map

| Hem. | Brain Region | BA | Coordinates |  |  | Vol. | T |
| --- | --- | --- | --- | --- | --- | --- | --- |
|  |  |  | (X, Y, Z) |  |  |  |  |
| L | Insula | 48 | -36 | 6 | 0 | 12 | 5.79 |
| R | Insula | 48 | 39 | 3 | 6 | 37 | 4.75 |
|  | Temporal pole: superior temporal gyrus | 38 | 51 | 6 | -9 |  | 4.12 |
|  | Insula | 48 | 42 | 9 | -3 |  | 4.09 |
| L | Insula | 48 | -45 | 6 | 9 | 10 | 4.54 |
|  | Insula | 48 | -42 | -3 | 9 |  | 3.82 |
| R | Median cingulate and paracingulate gyri | 32 | 3 | -9 | 42 | 16 | 4.12 |
|  | Median cingulate and paracingulate gyri | 23 | -3 | -21 | 45 |  | 4.07 |
| R | Cerebellum_Superior | 18 | 9 | -63 | -21 | 10 | 4.07 |

Note: all the results reported above were significant at $p < 0.001$ , $k > 10$ , uncorrected at the voxel level.

#### Task-dependent FC for predicting the third-party framing effect

As shown in Fig. 4D-E, regarding the brain network generated from the overlap between the T-contrast map and the regression map, the FC-based RVR models significantly predicted the third-party framing effect size (*r* = 0.349, permutation *p* = 0.031). The connections that showed strongest absolute weight (ļweightļ > 0.15) were: the FC between the left insula (MNI [−36, 6, 0]) and superior cerebellum (MNI [9, −63, −21]) (weight = 0.92), between the right insula (MNI [39, 3, 6]) and superior cerebellum (weight = 0.16), between the right insula and left insula (MNI [−45, 6, 9]) (weight = −0.15). These machine learning-based results remained significant under different FC thresholds, suggesting the robustness of our major findings.

#### ROI analysis

We selected the left insula (MNI [−36, 16, 2]) and cerebellum (MNI [18, −72,-24] for anger, MNI [−16 −70 −21] for disgust, and MNI [24 −82 −28] and [16 −84 −26] for mentalizing) as our ROIs. Paired sample *t*-tests revealed that the *harm* frame elicited significantly stronger activation than the *help* frame in the left insula (*p* = 0.009) and the ROI in the cerebellum for anger (*p* = 0.012) but not the ROI in the cerebellum for other functions (*ps* > 0.185) (Fig. 5).

**Figure 5.**
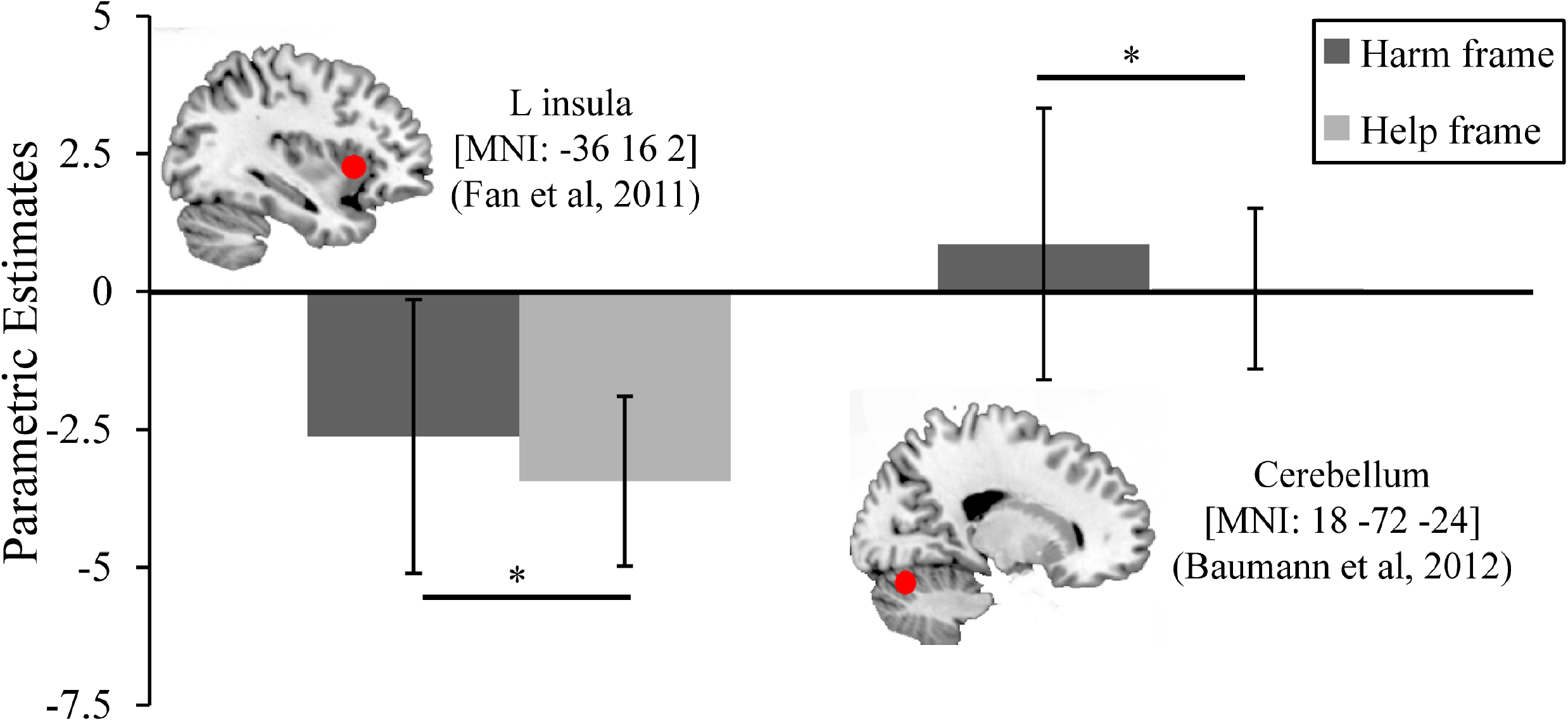
Results of region of interest analysis.

#### DCM

In the RVR analysis, the FC between the left insula (MNI [−36, 6, 0]) and superior cerebellum (MNI [9, −63, −21]) exhibited the strongest weight in relation to the third-party framing effect. We used DCM to further examine the direction of the FC between these two regions and how it was modulated by different frames. Results showed that the model defined by bidirectional modulation effect that worked on the bidirectional connection between the left insula and the superior cerebellum best accounted for the data (Fig. 6A, exceedance probability [xp] = 0.55, mean-variance explained 27.53%), while all the other models exhibited significantly lower probabilities (*xps* < 0.05; see details in Fig. 6B).

**Figure 6.**
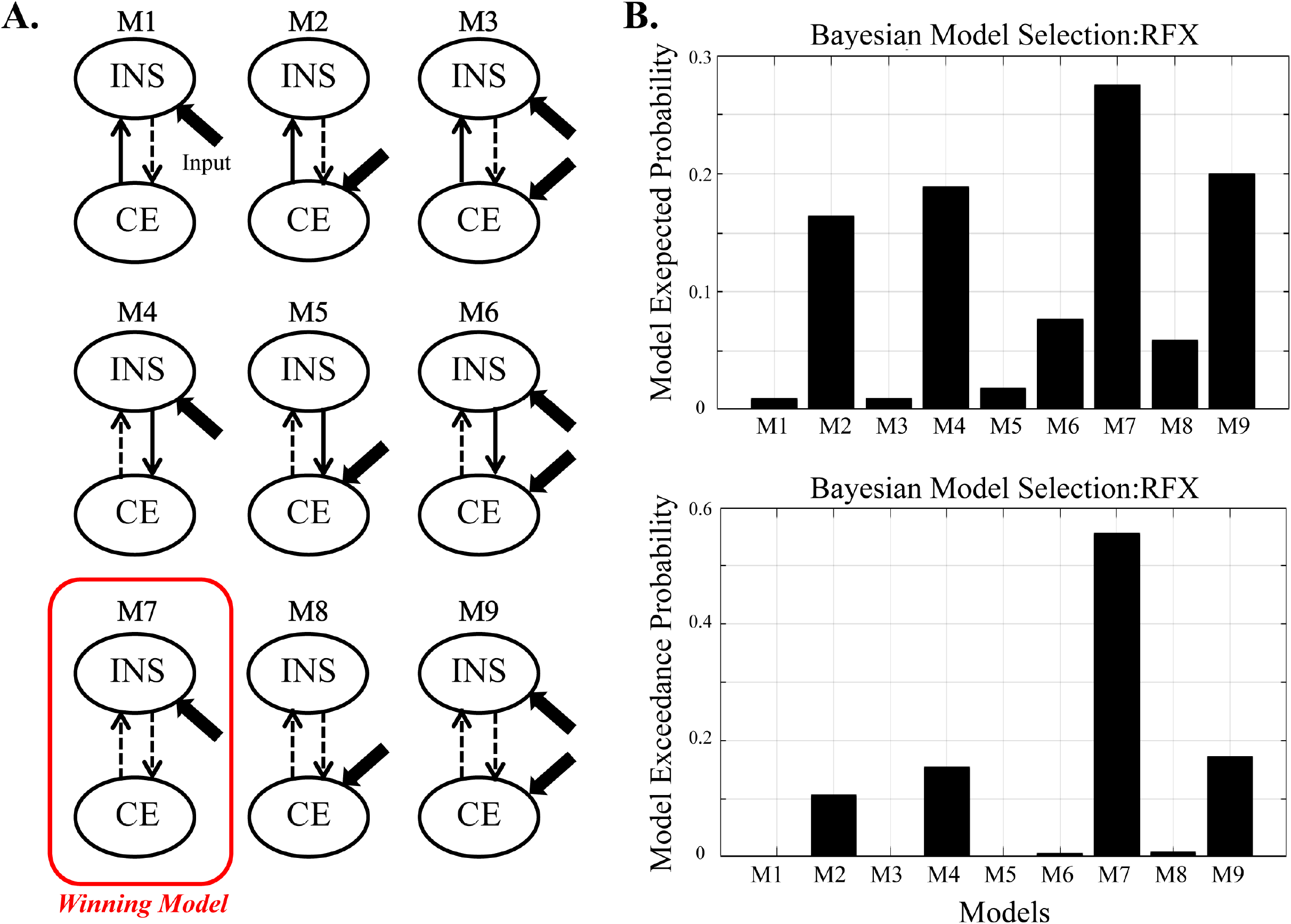
Results of dynamic causal modeling. (A) All possible nine models. The grey lines show where the modulation effect lied in. The winning model (M7) is marked with a red rectangle. (B) Results of Bayesian model selection.

## Discussion

In this series of experiments, we asked participants to observe two putative players interacted. Specifically, player A chose between “harming” and “not harming” player B in the *harm* frame condition, and between “helping” and “not helping” player B in the *help* frame condition. These two conditions were objectively but not verbally equivalent, conforming to the definition of frame manipulation (Rabin, 1998). In Experiment 1, we first examined whether third-party punishment could be modulated by the framing effect at the behavioral level. The results confirm our hypothesis, such that the participants were more willing to punish player A (at the cost of their own benefits) in the *harm* frame condition than in the *help* frame condition, resulting in a third-party framing effect. In our opinion, this effect reflected that “harming others” was judged by the participants as a more serious norm violation than “not helping others” (Crockett et al., 2014). The framing effect was robust unless player A was highly pro-helping, which is reasonable because it would be inappropriate to punish player A in this situation. We then calculated the averaged framing effect across different moral levels except “highly pro-helping” and found that it was significantly larger than zero. Likewise, self-reported “empathic feeling to player B” and “moral outrage to player A” were both larger in the *harm* frame condition than in the *help* frame condition. Moreover, these three behavioral indexes were significantly correlated between each other. To determine the relationship between these variables, we then conducted a mediation analysis and found that moral outrage acted as a full mediator between empathy and the third-party framing effect. The mediation effect indicates that compared to the *help* frame condition, enhanced empathic feelings to victims (player B) in the *harm* frame condition turned into stronger moral outrage toward norm violators (player A), and finally manifested as harsher punishment at the behavioral level (see also Shah et al., 2020).

In Experiment 2, we further combined our experimental paradigm with the fMRI technique. At the individual level, the whole-brain contrast comparing different frame conditions shows that the bilateral insula, MCC, and right cerebellum were activated stronger in the *harm* frame condition than in the *help* frame condition. At the group level, insula activation increased as a function of the framing effect size. The insula has been associated with various forms of prosocial behavior, including third-party punishments, allocating resources equally, and rejecting unfair distributions (Hsu et al., 2008; Sanfey et al., 2003; Stallen et al., 2018). Recently, Sellitto et al. (2020) found that higher insular activity indicates stronger sensitivity to other’s losses and more generous choices in a framing task. In light of the literature (e.g., Jabbi et al., 2007; Singer et al., 2004), we suggest that the empathic function of the insula is the key to explain its impact on decision-making. That is, empathic representation of other’s (current or predictive) state within the insula is integrated into behaviorally relevant computation and finally leads to more prosocial decisions (see also Singer et al., 2009). Critically, the insula belongs to a core network of empathy for other’s pain (Jackson et al., 2005; Laneri et al., 2017; Yao et al., 2016). In the *harm* frame condition of this study, participants observed that player A intentionally chose to “harm” player B; in this case, those who have stronger empathic responses (indexed by higher insular activation) to player B may feel that player A’s decision is a more serious norm violation and therefore deserves to be punished (see also Krueger & Hoffman, 2016).

Our findings about the cerebellum might be surprising, seeing that this brain area has been most often associated with movement-related functions (e.g., motor control) in the literature (Glickstein, 2007). However, Schmahmann and Caplan (2006) point out that it may no longer be appropriate to view the cerebellum as purely a motor control device; rather, accumulating evidence suggests that the cerebellum contributes to emotional processing and social cognition in many ways (Baumann & Mattingley, 2012; Turner et al., 2007; Van Overwalle et al., 2014). For instance, specific regions in the cerebellum are involved in encoding negative emotionally-evocative stimuli such as noxious heat and unpleasant images (Moulton et al., 2011). Most relevantly, the cerebellum subserves anger, aggression, and violence that are pertinent to moral judgments and actions (Demirtas-Tatlidede & Schmahmann, 2013). Regarding the versatile functions of the cerebellum in socio-emotional processing, our follow-up ROI analysis aimed to clarify whether cerebellum activation in this study could be accounted for by moral outrage (that is, the anger of observing moral violation: Landmann & Hess, 2017; Rothschild & Keefer, 2017). To do this, we selected an ROI within the cerebellum that corresponds to anger experience according to previous studies (Baumann & Mattingley, 2012); meanwhile, other ROIs that corresponds to disgusting experience and mentalizing were also selected for comparison (Baumann & Mattingley, 2012; Van Overwalle et al., 2014). The results of ROI analyses reveal that compared to the *help* frame condition, the ROI in the cerebellum related to anger (but not disgust or mentalizing) was activated more strongly in the *harm* frame condition, which is consistent with previous literature that emphasizes the importance of anger in moral behavior and altruistic punishment (e.g., Nelissen & Zeelenberg, 2009; Salerno & Peter-Hagene, 2013). Also, our finding supports the viewpoint that the cerebellum is necessary for full consideration of the neural basis of moral cognition (Ganis et al., 2003; Harada et al., 2009; Marchewka et al., 2012).

To explore whether (and how) the above brain areas constitute a network underlying the third-party framing effect, we run an RVR analysis and found that the effective connectivity between the left insula and cerebellum showed stronger absolute weight compared to other neural connections when predicting the behavioral framing effect size. We then conducted a DCM analysis on these two nodes, the results of which indicate that frame information was first entered into the left insula before transferring to the cerebellum. In our opinion, these findings are in line with the behavioral results of Experiment 1 that moral outrage toward wrongdoing (corresponding to the cerebellum) may mediate the relationship between empathy toward victims of norm violation (corresponding to the insula) and third-party punishment. Accordingly, a possible mechanism of third-party framing effect could be identified: our frame manipulation could trigger different levels of empathic response to victims by highlighting (or omitting) the severity of norm violation; this response indirectly drives altruistic punishment by modulating moral outrage toward norm violators. Here, the left-lateralized hemisphere asymmetry of the results might also have implications: while the right insula has been more frequently involved in an affective-perceptual form rather than a cognitive-evaluative form of empathy, the left insula is suggested to be recruited in both of these forms (Bird et al., 2010; Gu et al., 2013; Koban et al., 2013; Mazzola et al., 2010). Regarding that, we suggest that cognitive (but not affective) empathy was engaged in this study, possibly because our participants were unable to directly observe other people’s physical manifestations of pain (see also Cui et al., under review).

People punish norm deviation not only when they are the victims (i.e., second-party), but also – and more importantly – when they witness unrelated others being victimized (i.e., third-party) (Fehr & Fischbacher, 2004). Across societies, moral norms are enforced predominantly through third-party rather than second-party punishment (Bendor & Swistak, 2001; Buckholtz & Marois, 2012). Nonetheless, an act of norm violation may or may not be punished by third-parties, depending on various personal and situational factors (Kallgren et al., 2000; Sharma et al., 2014). In this study, we showed that third-party punishment could be strengthened when a choice is framed as intentionally harming others. As pointed out by Haidt and Graham (2007), evolutionary history has shaped maternal brains to be highly sensitive to signals of cruelty and harm; therefore, people generally try to prevent or relieve other’s harm, thus making the harm/care norm a strong moral restriction. Correspondingly, observing others violating this norm could provoke strong moral outrage that fuels costly punishment (Hartsough et al., 2020; Rothschild et al., 2013). In contrast, when a choice is framed as merely “not helping others,” that may be less likely to trigger the harm/care norm in people’s minds. Overall, this study reveal that third-party punishment is vulnerable to decision frames, which may help understand the flexibility of moral standards and moral actions (FeldmanHall et al., 2018). More broadly speaking, the current findings enrich the knowledge about the psychological processes of moral judgment and their corresponding neural underpinnings (Kelly & O’Connell, 2020).

On the other hand, our findings may broaden the understanding of the framing effect in general. According to previous studies, a stronger framing effect in non-social contexts (e.g., lottery) is associated with higher activation in the amygdala, possibly reflecting aversive emotional responses (e.g., anticipatory anxiety) triggered by heightened risk perception under frame manipulation (De Martino et al., 2006; Xu et al., 2013). Meanwhile, the framing effect in social contexts mainly employs the temporoparietal junction, which may reflect the process of perspective taking that grounds empathic emotions (Liu et al., 2020). To our knowledge, these studies have investigated the framing effect from a first-party perspective, that is to say, frame manipulation affects participants’ feelings about the decision outcomes that they would receive. In contrast, frame manipulation in this study modulates vicariously activated moral emotions based on third-party observation (see also Hartsough et al., 2020) and therefore recruits distinct neural circuits compared to that underlying the classical framing effect. Still, our findings support the idea that the framing effect is essentially an emotional phenomenon regardless of whether it is based on first-party or third-party experience.

Below we propose some future directions for follow-up studies to consider. First, frame manipulation could be conducted on different aspects (e.g., reference point vs. outcome salience) of a decision dilemma (Kühberger, 1998). While our task focuses on outcome salience, it would be interesting to see whether the third-party framing effect based on reference point manipulation would show different mechanisms. Second, aside from punishing norm violators, another important kind of third-party intervention is to compensate the victims (Leliveld et al., 2012). Investigating the framing effect on third-party compensation would have important implications for promoting prosocial behavior. Finally, one of the major threats to altruistic third-party punishment is “diffusion of responsibility,” that is, the presence of bystanders may reduce an unaffected observer’s likelihood to take actions (Darley & Latane, 1968). Relevantly, one of our studies showed that diffusion of responsibility inhibits neural activation in the insula (Feng et al., 2016). Researchers may examine whether appropriately using frame manipulation would counteract the negative influence of diffusion of responsibility.

## Author Contributions

The first three authors (#) contribute equally. F.C. designed the research; J.Y., K.D., and X.H. conducted the experiments and collected data; J.L. and J.Y. analyzed data; F.C., J.L., R.G., and J.Y. wrote the paper.

## Acknowledgment

This study was funded by the National Natural Science Foundation of China (31871109 to F.C., 31900779 to J.L., and 32071083 to R.G).

## Ethical standards

All procedures performed in this study were in accordance with the 1964 Helsinki declaration and its later amendments or comparable ethical standards. The local ethics committee approved the experimental protocol.

